# Membrane-proximal external region is a superior target for mediating effector activity of HIV-1 specific chimeric antigen receptor modified T cells

**DOI:** 10.1101/2020.03.11.987610

**Authors:** Emiko Kranz, Joshua Chan, Maya Hashimoto, Toshio Kanazawa, Hanlu Wang, Masakazu Kamata

## Abstract

The use of chimeric antigen receptor modified-T (CAR-T) cells in adoptive immunotherapy has been popularized through recent success in the field of cancer treatment research. CD4ζ CAR, which targets HIV-1-infected cells, has been developed and evaluated in patients. Though well-tolerated for over a decade, efficacy was disappointingly limited. This result encourages us to develop a novel CAR more effective than CD4ζ CAR. To quantitatively compare anti-HIV-1 activity of different CAR constructs in a highly sensitive and reproducible manner, we developed a multicolor flow cytometry method for assessing anti-HIV-1 effector T-cell activity. “Target” Jurkat cells inducibly expressing an HIV-1_HXBC2_ envelope protein and “Non-target” control cells were genetically labeled with red and blue fluorescent protein, respectively, and co-incubated with human primary T cells transduced with anti-HIV-1 “Effector” CARs at various Effector vs Target cell ratios. Absolute cell numbers of each population were collected by MACSQuant Analyzer and used for calculation of relative cytotoxicity. We successfully ranked the cytotoxicity of three previously reported single chain-antibody CARs and six newly developed single-domain antibody CARs in comparison to CD4ζ CAR. Interestingly, three CARs—10E8, 2E7, and 2H10—which demonstrate high cytotoxic activity were all known to target the membrane-proximal external region. Use of this novel assay will simplify assessment of new CAR constructs and in turn accelerate the development of new effective CARs against HIV-1.

**Author Summary:** Adoptive immunotherapies that utilize autologous T cells expressing a desired antigen-specific CAR aim to elicit directed immune responses. In recent years, CAR immunotherapies have been promoted extensively in B cell malignancy treatments. The HIV-1-targeting CAR, known as CD4ζ, was developed over 20 years ago and has been widely and longitudinally tested in patients. However, its effectiveness was hindered by poor survival and functionality of the transduced cells. To conduct quantitative evaluation of newly designed anti-HIV-1 CARs, we developed a novel multicolor flow-based assay for HIV-1-specific cytotoxicity, enabling sensitive and quantitative assessment in a high-throughput fashion. This assay would be also useful in screening HIV-1-targeting immune receptors—including CARs and T cell receptors—and other immunotherapeutic drugs such as anti-HIV-1 antibodies.

## Introduction

Chimeric antigen receptors (CARs) are artificially engineered receptors that confer a desired specificity to immune effector T cells such as CD4+ (as a helper T cell) and CD8+ (as a cytotoxic T cell) [1-6]. When a CAR encounters its target ligand, it signals the cell in a T cell receptor (TCR)-like manner, but not in a human leukocyte antigen (HLA)- dependent manner; thus, this approach can be utilized in treatments for anyone. The use of CAR-modified T (CAR-T) cells has been applied extensively in anti-cancer research, such as with solid organ tumors and lymphomas [6-12].

A promising candidate for use in anti-HIV-1 adoptive therapy is the CD4ζ CAR, which contains extracellular domains from CD4, a major HIV-1 receptor, and an internal signaling domain derived from CD247, a CD3ζ-chain. CD4ζ CAR utilizes the CD4 recognition site to respond to an HIV-1 envelope protein that lies on the exposed surface of infected cells. Once activated, the ζ-chain emits a signal to trigger potent effector function against infected cells [13-21]. This CAR has been widely and longitudinally tested in patients over 500 patient years [22-25]. Treatment was safe and well-tolerated for over a decade, but antiviral effects were limited, most likely due to poor maintenance of gene-modified cells. These results facilitated the restructuring of CD4ζ CAR to preserve its inherent potential as well as to heighten its antiviral capabilities, leading to the creation of a new CAR line: novel anti-HIV-1 CARs using a single chain (scFv) form of broadly neutralizing antibodies (bNAbs) [26-29].

Cytotoxic assays are used to determine the efficiency of a CAR’s cytotoxic activity by observing its ability to kill “Target” cells that express the epitope recognized by a corresponding “Effector” CAR. The radioactive chromium (^51^Cr)-release method developed in 1968 has traditionally been used to determine the cytotoxic activity of effector cells[30]. Although the assay is reliable and has become a “gold standard,” it has a number of disadvantages and functional limitations such as low sensitivity, risk of radioactive contamination, and spontaneous release of ^51^Cr contributing to heightened background levels that tend to limit its application in quantitative assessment and high-throughput screening. To overcome these issues, several nonradioactive methods have been developed that mainly use fluorescent dyes [31-42]. However, despite the advantage of working with nonradioactive material, these methods have not yet found a broad acceptance—likely due to the labor-intensive procedure, wide variability across assays, and low reproducibility in results.

To perform quantitative evaluation of newly designed CARs with high reproducibility and accuracy, we developed a novel multicolor flow cytometry-based assay for HIV-1-specific effector activity. The assay uses two cell lines: “Target”, which is genetically labeled by a red-fluorescent protein (mCherry) and inducibly expresses HIV-1_HXBC2_ envelope proteins, and “Non-target,” which is labeled by a blue fluorescent protein (TagBFP) but does not express the HIV-1 envelope proteins. To minimize non-specific cell death by mismatched major-histocompatibility complex (MHC), the surface expression of MHC was first eliminated from both Non-target and target cells by knocking out the β2 microglobulin (β2MG) gene with CRISPR gene editing technology[43]. In addition, low CD4-expressing populations of these cells were further selected to minimize spontaneous cell death initiated by HIV-1 envelope mediating cell fusion[44]. Equal numbers of Target and Non-target cells are co-incubated with CAR-T effector cells in various ratios for 16 hours. HIV-1-specific cytotoxicity is determined by counting absolute cell numbers of each population. This assay enables assessment of HIV-1-specific cytotoxicity at a single cell level, allowing for quantitative and simple numerical analysis of results rather than relying on measurements of a released substance. These benefits expand the applicability of the assay, allowing for personalization according to the samples present without sacrificing efficiency.

## Results

### Generation of target cells for HIV-1-specific effector cell assay

The ^51^Cr-release assay has been known as the gold standard for evaluating effector T cell activity [30]. The main difficulty with the ^51^Cr assay lies in the complications that arise from variation in ^51^Cr labeling due to spontaneous release of ^51^Cr, elevating background levels substantially. The success and analytical quality of flow cytometry-based assays hinges considerably on the reproducibility of results—the basis of this consistency lies in the quality of an assay’s Target cell population. To remedy these issues presented by the ^51^Cr assay, we have developed a novel cell line stably expressing fluorescent protein that is not spontaneously released, unless the integrity of the cell surface membrane is compromised due to attacks by such as cytotoxic T cells. We utilized two different Jurkat cell lines previously established: Target, which inducibly expresses HIV-1_HXBC2_ envelope proteins upon doxycycline (DOX) removal from the culture medium, and Non-target, which does not express the HIV-1 envelope proteins [45]. To minimize cytotoxicity mediated by allogeneic reactions because of mismatched MHC, we first eliminated MHC surface expression via gene targeting of β2MG using CRISPR/Cas9 technology (**S1 Fig**). The cells were then modified by genetic labeling with mCherry for Target cells (HXBC2) or TagBFP for Non-target cell (ΔKS). Fluorescently labeled populations with missing HLA expression were sorted by FACSAria II along with a CD4 dimmer population to minimize cell fusion induced by the interaction between HIV-1 envelope and CD4 (**Figs 1B,C)**. HIV-1 envelope protein expression of resultant cells was confirmed by western blotting using antibodies against HIV-1 gp120 (2G12) and gp41 (2F5) 4 days after removal of doxycycline (DOX) from the culture medium (**Fig 1D**) as well as flow cytometry using a fusion protein of soluble human CD4 and Fc portion of human IgG1 (sCD4 Fc), followed by APC-conjugated anti-human Ig Fc (**Fig 1E**). Importantly, levels of HIV-1 envelope expression in HXBC2 on days 9 and 12 were similar to those of primary human CD4 T cells infected by two different HIV-1 strains.

**Fig 1.**
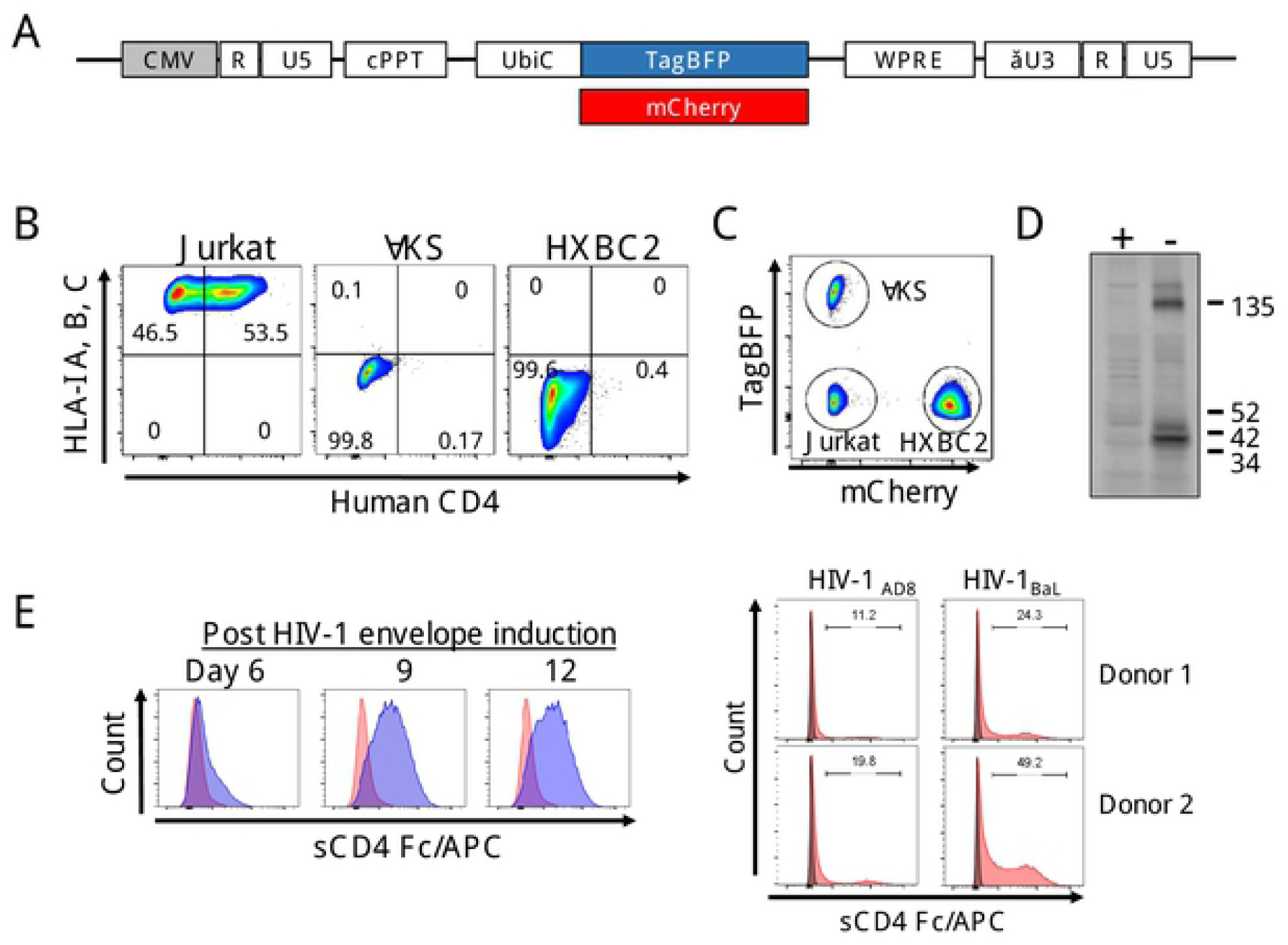
Generation of Jurkat cells inducibly expressing envelope protein from HIV-1_HXBC2_ (HXBC2). **(A)** Schematic of lentiviral vectors for expressing TagBFP and mCherry. These vectors have an FG12-derived backbone possessing a self-inactivating LTR, central polypurine tract (cPPT), ubiquitin C promoter (UbiC), and a mutant Woodchuck Hepatitis Virus Posttranscriptional Regulatory Element (WPRE). **(B)** Jurkat cells with or without inducibly expressing envelope protein from HIV-1_HXBC2_ first had human-β2 microglobulin (β2MG) expression knocked out by CRISPR-Cas9 gene editing using a non-integrating lentiviral vector encoding Cas9 together with sgRNA specific for β2MG as shown in **S1 Fig**. Populations missing human-leukocyte antigen (HLA)-1 A, B, and C expression were negatively enriched by magnetic bead separation, followed by transduction with a lentiviral vector encoding either TagBFP for ΔKS or mCherry for HXBC2. Cells were then labeled by PE/Cy7-conjugated anti-human HLA-I A,B,C and BV711-conjugated human CD4 antibodies then further selected for TagBFP^+^/HLA-I A,B,C^-^/CD4^dim^ (ΔKS) or mCherry^+^/HLA-I A,B,C^-^/CD4^dim^ (HXBC2) populations by FACSAria II flow sorter. **(C)** Equal numbers of ΔKS, HXBC2, and unmodified Jurkat cells were mixed and analyzed on LSRFortessa for TagBFP and mCherry expression. **(D)** HXBC2 cells were cultured for 4 days in the presence or absence of 1 µg/ml doxycycline (DOX) to induce HIV-1_HXBC2_ envelope expression. Five million cells were lysed in 1% CHAPS and HIV-1 envelope expression was analyzed by western blotting using a mixture of anti-HIV-1 gp41 (2F5) and anti-HIV-1 gp120 (2G12) antibodies. Numbers to the right of the picture indicate molecular mass in kilodaltons. +: with DOX, -: without DOX. **(E)** A fusion protein of soluble human CD4 and Fc portion of human IgG1 (sCD4 Fc) was used for detection of HIV-1 envelope expression on cell surface. One million HXBC2 cells cultured in the absence (red) or presence (blue) of DOX for 6, 9, and 12 days were incubated with 1 µg sCD4 Fc on ice, followed by APC-conjugated anti-human Ig Fc portion. Human primary T cells (two donors) were infected with HIV-1_AD8_ and HIV-1_BaL_ and used as a positive control for levels of cell surface expression ofHIV-1 envelope proteins detected by sCD4 Fc. Gray: uninfected human primary T cells, Red: HIV-1-infected cells.

### Establishing HIV-1-specific cytotoxicity assay using multicolor flow cytometry

Assessment of cytotoxicity in our assay is carried out by counting absolute live cell populations in each well following co-culture of “Effector”, “Target”, and “Non-target” cells (**Fig 2**). Target and non-target cells that have been genetically modified to express different fluorescent proteins are easily characterized by multicolor flow cytometry. Upon attack by effector cells, their cytoplasmic contents are released into the supernatant and the number of fluorescent cells decreases, i.e., the loss of fluorescence from cells is used as an indicator of both decreased membrane integrity and cell death [46]. The same numbers (10,000 cells/well in a 96 well plate) of HIV-1 envelope-expressing target cells are co-cultured with non-target cells lacking envelope expression and effector cells designed to eliminate target cells (**Fig 2A**). Isolated populations of target cells are counted against those of non-target cells to evaluate apparent cytotoxic activity following co-culturing with effector cells (**Figs 2B,C**). Our assay is unique not only in its use of uniform fluorescent marking, but also in its utilization of the MACSQuant system for counting actual live cell populations. The greatest advantage of this instrument is the fully automated system, which allows the collection of data in a high throughput manner with easy operation. Use of automated counting by MACSQuant also proves advantageous in its consistency, removing complications by human error.

**Fig 2.**
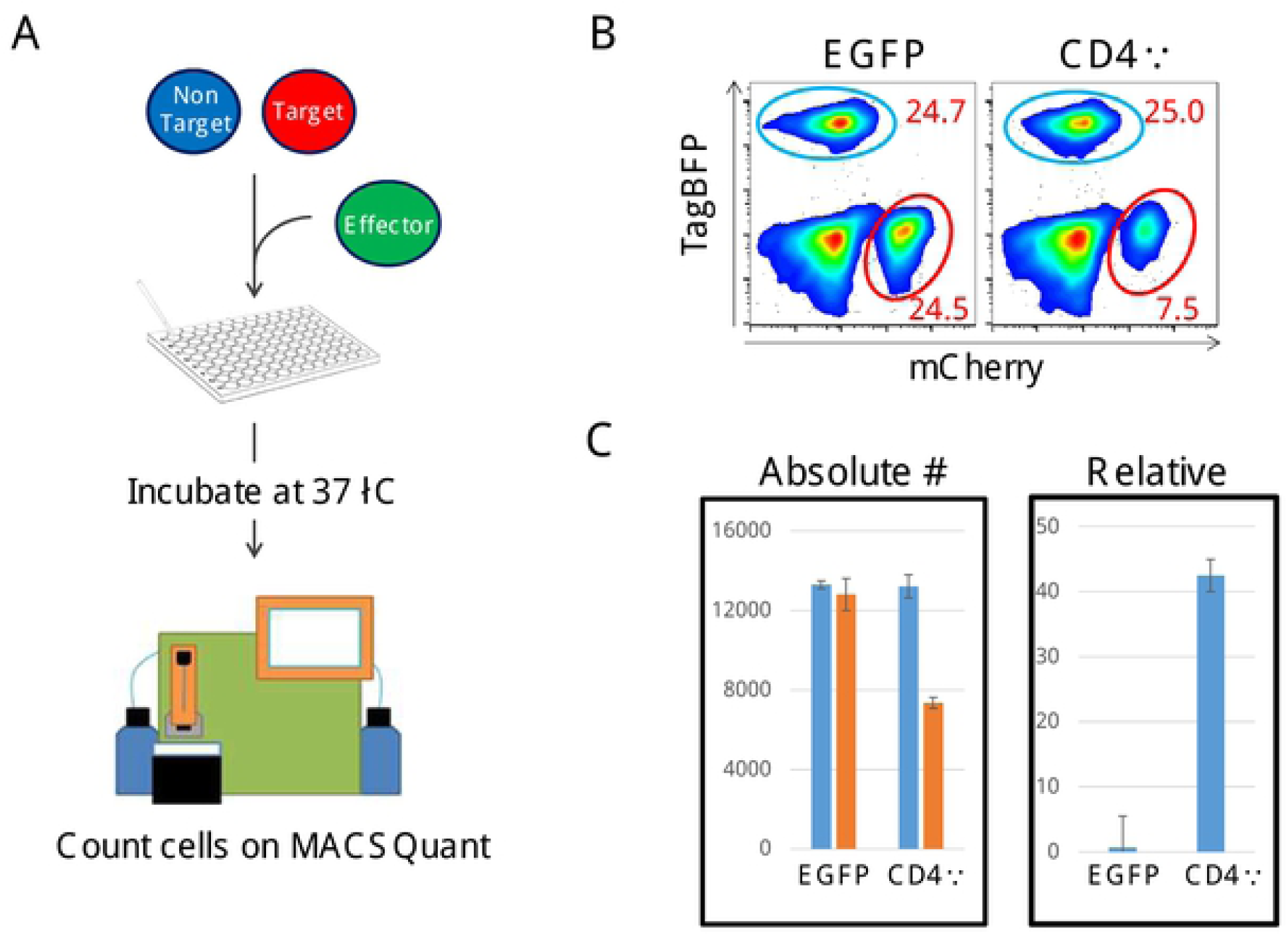
Schematic outline of a multicolor flow cytometry-based cytotoxicity assay designed for HIV-1 specific CAR-T cells. **(A)** Scheme of fluorescent-based cytotoxicity assay. TagBFP^+^ control Jurkat cells (ΔKS; Non Target) and mCherry^+^ Jurkat cells that inducibly express HIV-1_HXBC2_ envelope protein upon removal of DOX from culture medium (HXBC2; Target). ΔKS and HXBC2 cells were co-incubated together with human primary T cells transduced with a vector encoding CD4ζ CAR or a control EGFP vector (CD4ζ or EGFP). (**B,C**) Absolute cell number of each population circled in blue (Non-target) or red (Target) was analyzed by MACSQuant. HIV-specific cytotoxicity was calculated as relative cytotoxicity as follows: Relative cytotoxicity (%) = 100 x (1 - Target cell numbers/Non-Target cell numbers). The uncircled population is composed of effector cells. Data represent the mean ± standard deviation from triplicate wells.

### The assay shows high reproducibility with wide dynamic range

A comparative assay system relies heavily on consistency of results; such consistency may be assured by adjusting variables that the cytotoxic effect depends on. Before we use the above established cell lines for further assays, cells were processed to isolate single-clone populations since bulk populations consist of heterogeneous cells that affect the cytotoxicity assay (**Fig 3**). Based on levels of HIV-1 envelope expression and sensitivity to CD4ζ CAR-mediated cytotoxicity, we selected HXBC2 clone #39 and ΔKS clone #13 for later experiments. Clone #39 showed stable HIV-1 envelope expression over 7 days after 8 to 10 days of culture with medium containing no DOX (**Fig 3B**) and was able to mediate HIV-1-specific T cell proliferation (**Fig 3C**). HIV-1 specific cytotoxicity was confirmed after 8 hours of incubation with CD4ζ CAR-expressing T cells and plateaued after 16 hours of incubation (**Fig 3D**).

**Fig 3.**
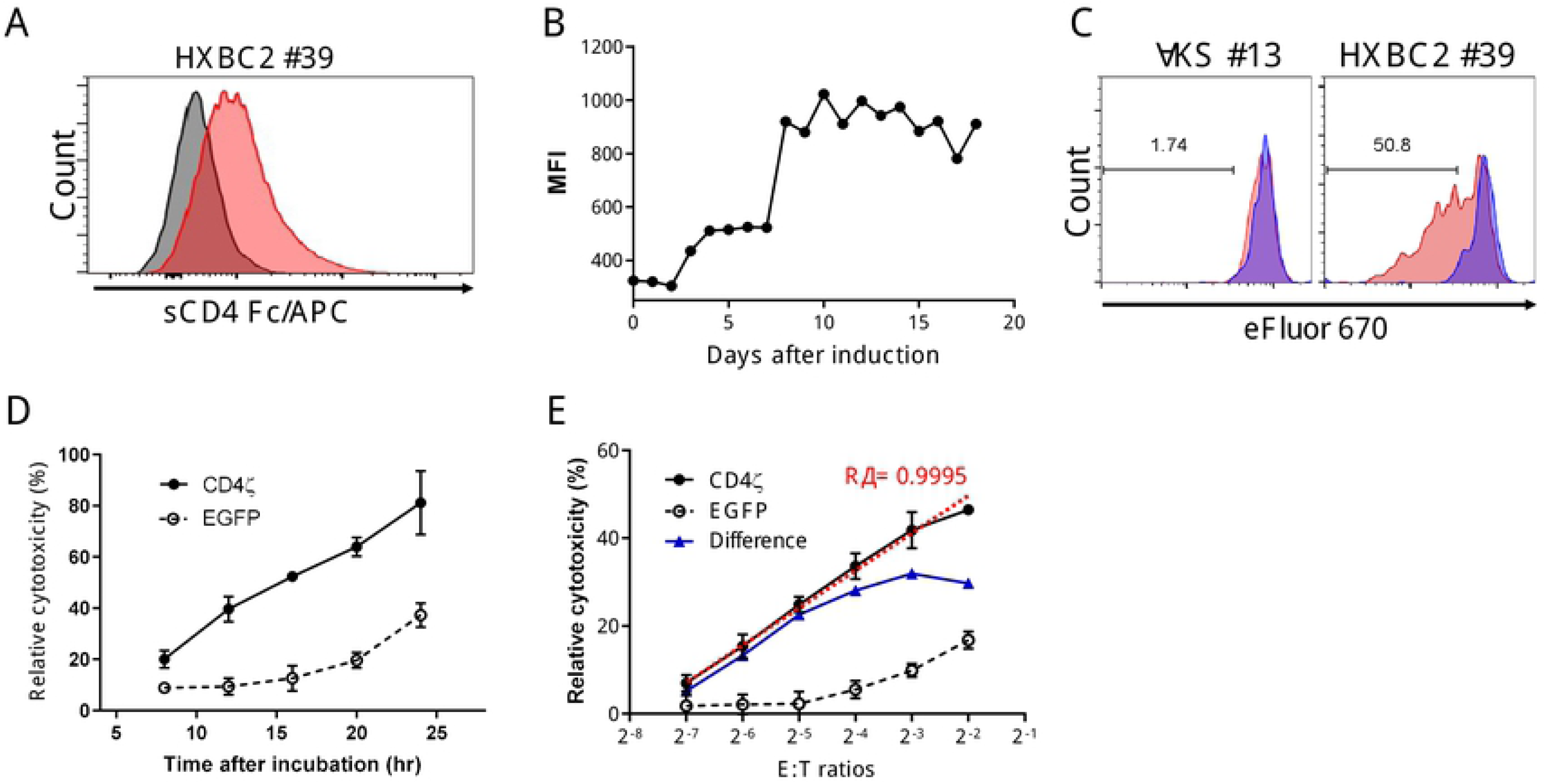
Characterization of single cell-derived HXBC2 cells. Single cell-derived clone for Target (HXBC2 #39) and Non-Target (ΔKS #13) cells was obtained by limiting dilution. **(A)** HXBC2 #39 cells were cultured in medium with no DOX for 10 days. HIV-1 envelope expression was monitored by sCD4 Fc/APC staining as described in **Figure 1** (red). Uninduced HXBC2 #39 cells were used as a negative control (gray). **(B)** Surface expression of HIV-1_HXBC2_ envelope protein was monitored daily over 18 days and mean fluorescent intensities (MFIs) were plotted. **(C)** CD4ζ CAR-expressing human primary T cells were co-cultured with ΔKS #13 or HXBC2 #39 (E:T ratio = 1:10) after labeling with cell proliferation dye eFluor 670. Proliferation of CD4ζ CAR-T cells was monitored by dye dilution after 6 days culture. (**D,E**) HIV-1-specific cytotoxicity mediated by CD4ζ CAR was tested with increasing incubation time (8 - 24 hours) **(D**) or different effector:target ratio (E:T ratios = 2^-7^ to 2^-2^) (**E**). Results were calculated as a relative cytotoxicity as described in **Fig 2** and shown by the mean ± standard deviation from triplicate wells.

The critical factor to take into consideration for a reproducible assay is the Effector to Target (E:T) ratio. If there are too few effector cells, cytotoxic activity will appear inefficient and produce inconclusive results; however, if there are too many effector cells, non-target cells are at risk of collateral cytotoxic effect, resulting in high background signals. This saturation of effector cells may also result in higher levels of cytotoxic effect that do not reveal any further understanding of the tested CAR’s efficacy. Should such excessive amounts of effector be used, one would see high levels of killing no matter the actual capabilities of the CAR. Thus, choosing a ratio before effector function plateaus is ideal. Ratios ranged from 2^-7^ to 2^-2^ (0.0078125 to 0.25) and levels of cytotoxicity by CD4ζ increased in a linear fashion following increased E:T ratio (**Fig 3E**). Although these levels also elevated in controls (**Fig 3E, EGFP**), the differences of relative cytotoxicity between CD4ζ and EGFP control increased in a linear fashion from 2^-7^ to 2^-3^ and reached a plateau (**Fig 3E, Difference**). We therefore used the E:T ratios from 2^-7^ to 2^-3^ thereafter.

### Anti-HIV-1 CARs targeting gp41 MPER exert a potent anti-HIV-1 specific cytotoxicity

Anti-HIV-1 antibodies neutralizing wide-spectrum of HIV-1 strains called broadly-neutralizaing antibodies (bNAbs) have been developed or isolated from HIV-1-infected individuals. These antibodies are known to recognize diverse epitopes on HIV-1 envelope proteins to differing extents and are utilized for developing a variety of anti-HIV-1 CARs [47]. Prior to developing novel anti-HIV-1 CARs using bNAbs, we first tested whether previously reported anti-HIV-1 bNAbs can recognize HIV-1 envelope proteins expressed on cells (**Fig 4**).

**Fig 4.**
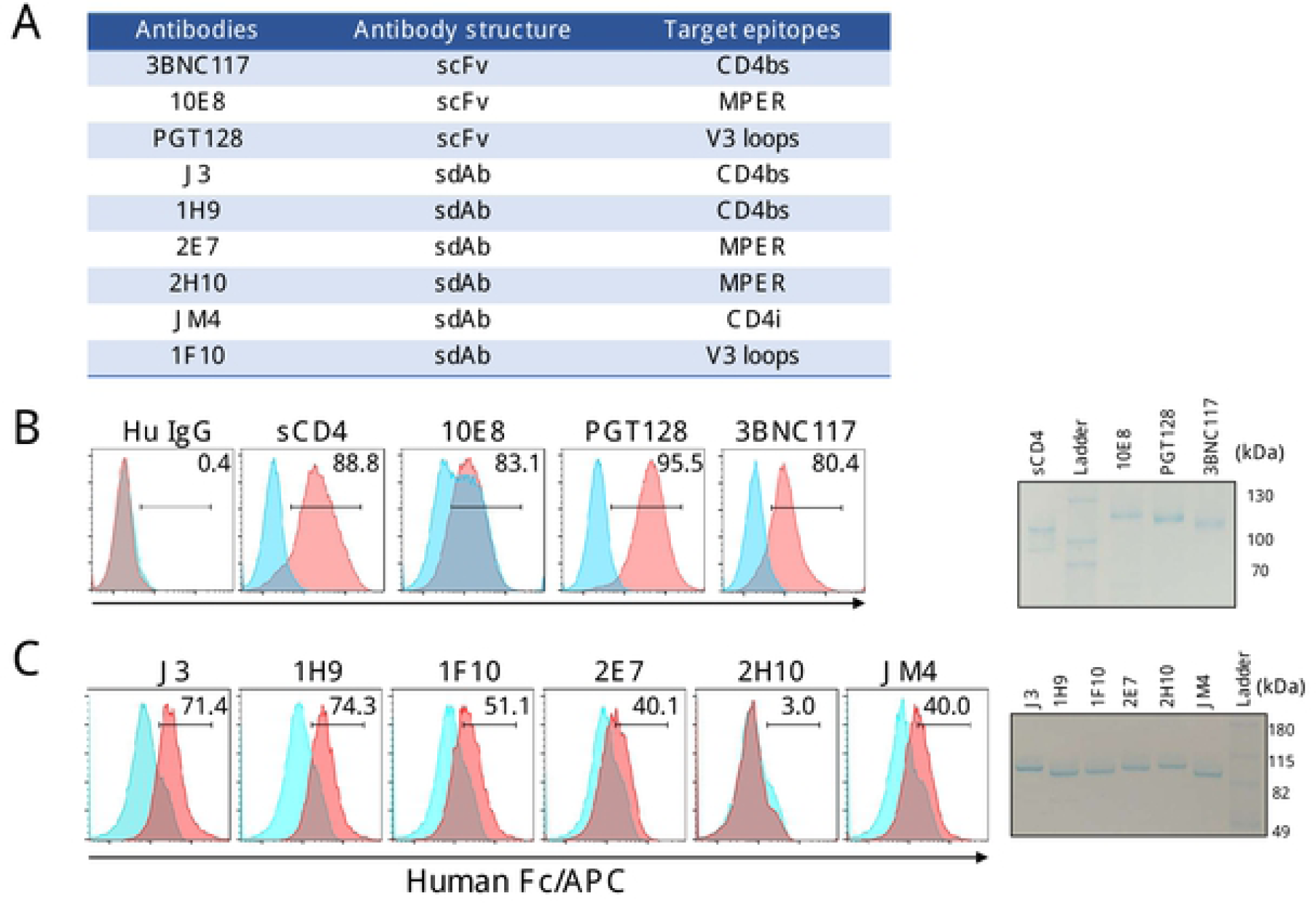
Binding features of anti-HIV-1 broadly neutralizing antibodies (bNAbs) to HIV-1_HXBC2_ envelope proteins expressed on HXBC2 #39 clone cells. **(A)** A list of anti-HIV-1 bNAbs used in this experiment. Recombinant forms of anti-HIV-1 bNAbs were expressed in 293T cells as a fusion protein with the Fc portion of human IgG1 and purified by protein A column. scFv: single-chain variable fragment. sdAb: single-domain antibody. **(B,C)** HXBC2 #39 cells were cultured in the absence of DOX for 10 days to induce HIV-1_HXBC2_ envelope protein. One million cells were incubated with 1 µg of each antibody on ice for 1 hour followed by APC-conjugated anti-human IgG Fc portion (red). Soluble CD4 Fc (sCD4) was used as a positive control and human IgG was used as a negative control. Uninduced HXBC2 #39 cells cultured in the presence of DOX were used to monitor non-specific antibody binding (blue).

We selected 10 different bNAbs based on their efficacies as well as a broad spectrum, including two different forms of antibody, conventional antibody and heavy-chain antibody [48, 49]. For CAR designing, the single-chain variable fragment (scFv) form or single-domain antibody (sdAb) form of these antibodies have been used.. Therefore we developed the scFV or sdAb forms of these antibodies and assessed whether they can recognize HIV-1 envelope proteins expressed on cell surface by flow cytometry. A portion of epitope receptor of scFv or sdAb form was conjugated to a common human immunoglobulin G1 (IgG1) Fc domain—termed synthetic-antibody mimetics (SyAMs)—allowing for quantitative immunological assays. All SyAMs can be enriched at >95% purity, as determined by SDS-PAGE gel, using a protein-A column (**Figs 4B,C**). HXBC2 #39 clone induced with envelope expression for 10 days were used for assessing the levels of epitope detection by those anti-HIV-1 SyAMs. Although epitope recognition differed between SyAMs, most were able to detect HXBC2 cells expressing HIV-1_HXBC2_ envelope proteins.

Three scFv and six sdAb fragments were then subcloned into the CD4ζ vector by replacement of the EGFP-P2A-CD4 fragment (**Fig 5A**). To improve CAR-T cell survival and their effector activities, all these CARs, including CD4ζ, included the 41BB costimulatory signaling domain inserted between the CD8 transmembrane domain and ζ-chain [50, 51]. To protect CAR-T cells from HIV-1 infection as well as virus production from proviral genes, all vectors included two anti-HIV-1 genes, C46 fusion inhibitor [52, 53] and shRNA against the LTR R region (sh516) as reported previously [54, 55]. The C46 gene was selected as the best inhibitor for HIV-1 infection in comparison with three other inhibitors: shRNA against CCR5 (sh1005) [56, 57], V2o[58], and AP3[59] (**S2 Fig**). All of these CARs were successfully expressed in human primary T cells at similar levels (**Fig 5B**).

**Fig 5.**
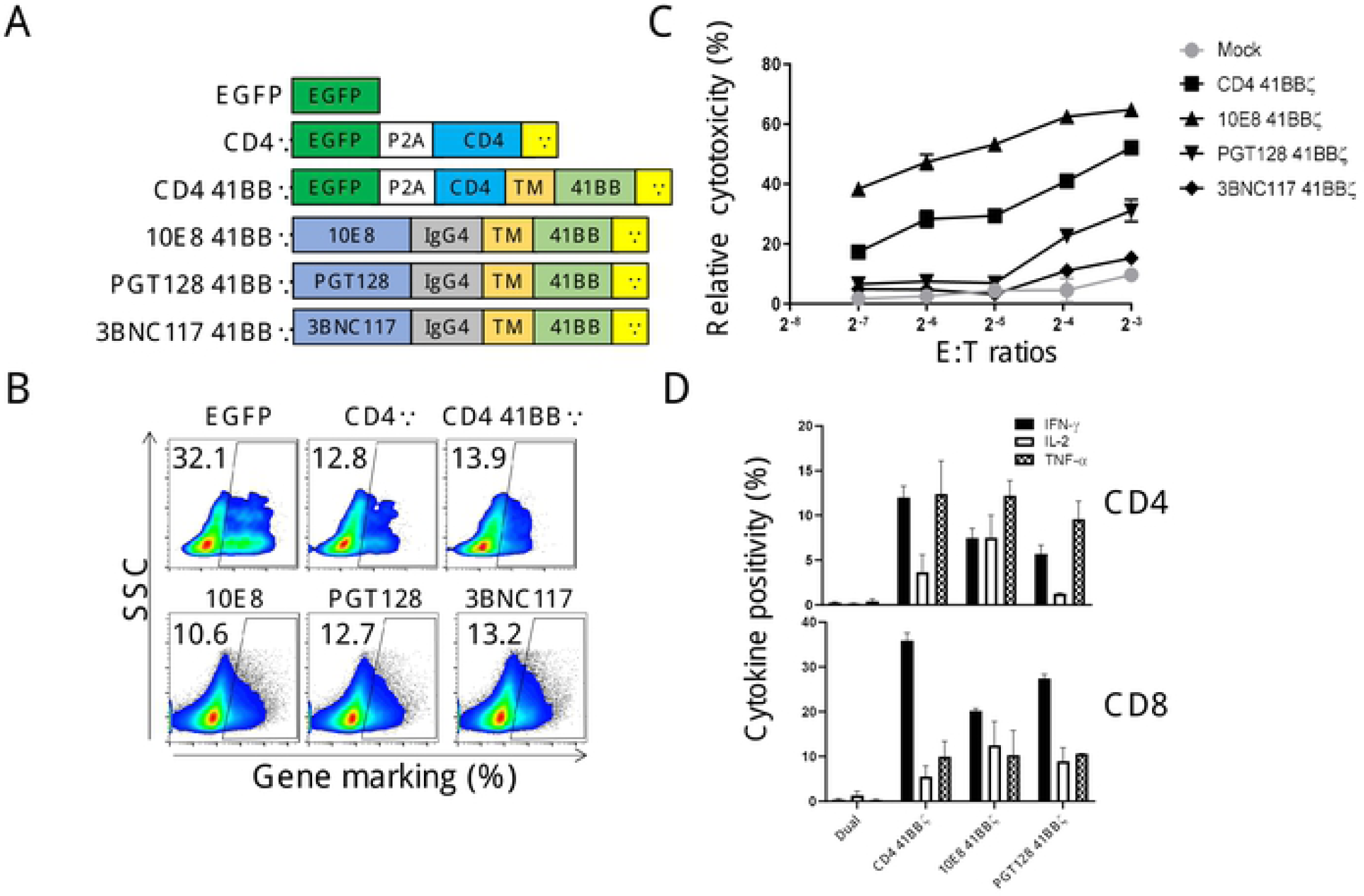
HIV-1-specific cytotoxicity mediated by anti-HIV-1 scFv CAR-T cells. **(A)** Schematic of anti-HIV-1 CAR structures expressed in lentiviral vectors. EGFP and CD4ζ serve as negative and positive controls. CD4 41BBζ CAR contains 4-1BB domain derived from human CD137. Three other CARs—10E8 41BBζ, PGT128 41BBζ, and 3BNC117 41BBζ—encode single-chain forms of anti-HIV-1 bNAbs. **(B)** Transduction levels of each anti-HIV-1 CAR were monitored by flow cytometry. EGFP, CD4ζ, or CD4 41BBζ transduction were monitored by EGFP expression. Three anti-HIV-1 scFv CARs were detected by staining with Alexa488-conjugated anti-human Fc portion. **(C)** Comparison of the specific cytotoxicity induced by anti-HIV-1 CAR. Data represent the mean ± standard deviation from triplicate wells. **(D)** Production of three different cytokines (IFN-γ, IL-2, and TNF-α) in CAR-modified CD4 and CD8 T cells was monitored by flow cytometry. Data were shown by % positivity of cytokine-producing cells.

Anti-HIV-1-specific effector activity of each CAR was ranked by a cytotoxicity assay developed above using CD4 41BBζ CAR as a reference CAR. To input the same number of effector cells in the assay, percent positivity of effector cells in each well was adjusted by unmodified mock T cells. 10E8 41BBζ CAR showed near 2-fold higher activity than that of CD4 41BBζ CAR. PGT128 and 3BNC117 41BBζ CARs showed a weak to no detectable cytotoxicity. 10E8 and PGT128 41BBζ CARs were further evaluated for cytokine production (**Fig 5D**). Compared to the no-CAR control (**Fig 5A, EGFP**), CD4+ and CD8+ T cells modified with these CARs successfully produced cytokines in response to induced HXBC2 cells expressing HIV-1 envelope proteins. Interestingly, there was no clear correlation between levels of cytokine production and induction of cytotoxicity; PGT128 CAR showed weaker effector activity than 10E8 CAR, but similar levels of IFN-γ and TNF-α, both are well-known pro-inflammatory cytokines [60], production in both CD4 and CD8 T cells.

Lastly we tested sdAb-based CARs in the same way. Overall, slightly lower levels of gene marking were seen with these CARs (**Fig 6B**), probably due to the use of different assay methods between scFv and sdAb CARs (see **Materials and Methods**). As seen with 10E8 41BBζ CAR, both MPER-specific sdAb-based CARs, 2H10 and 2E7, showed around 2-fold higher activity in comparison to that of CD4 41BBζ CAR. Except J3 41BBζ CAR, the three other CARs (1F10, JM4, and 1H9) exhibited similar levels of effector activity in comparison to that of CD4 41BBζ CAR (**Figs 6C,D**).

**Fig 6.**
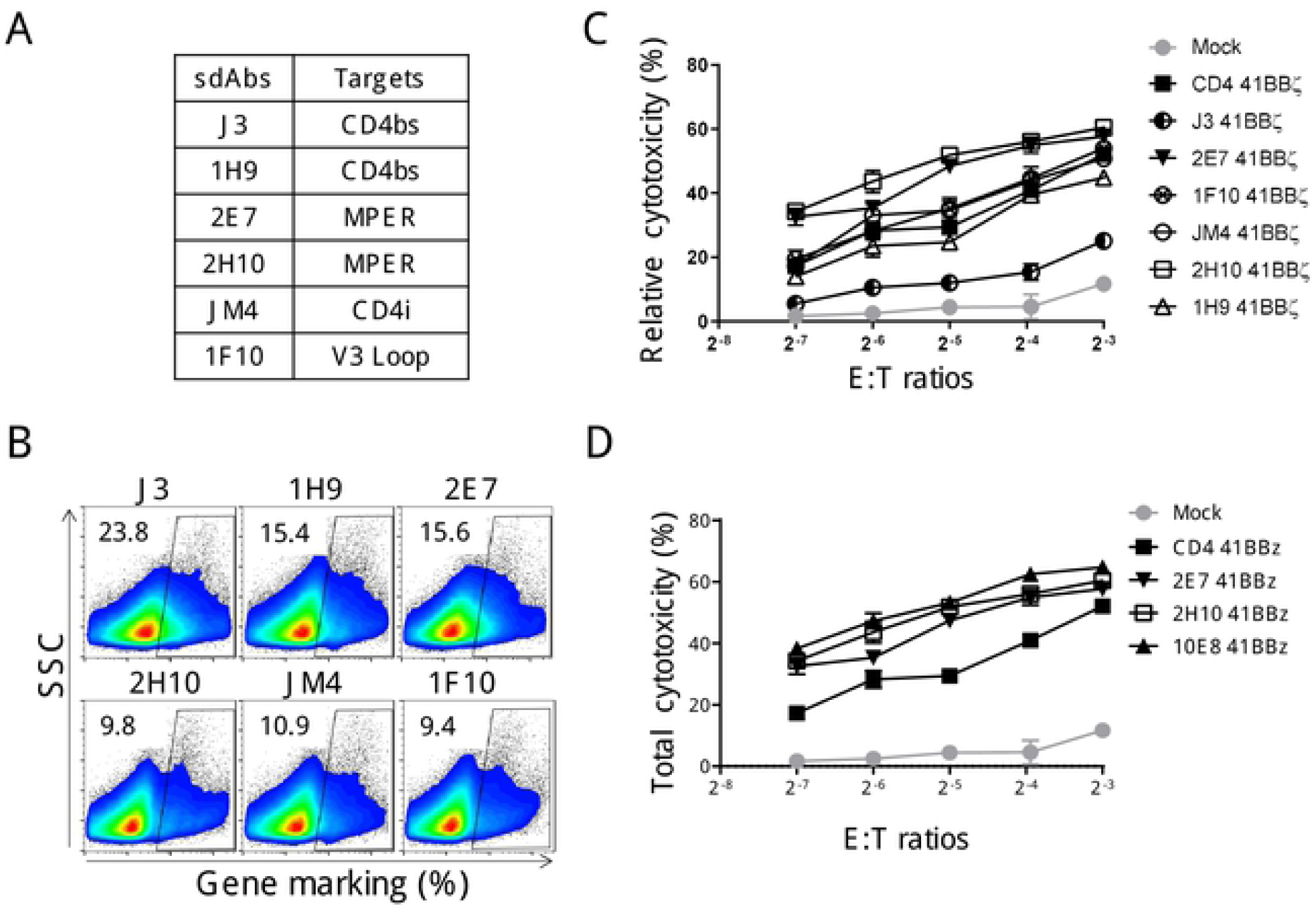
Superior anti-HIV-1 activity mediated by MPER-targeting anti-HIV-1 CAR. **(A)** A list of anti-HIV-1 sdAbs used for anti-HIV-1 CAR design. **(B)** Transduction levels of each anti-HIV-1 sdAb CAR were monitored by flow cytometry. Surface expression of anti-HIV-1 sdAb CARs was detected by biotinylated-protein L, followed by APC-conjugated streptavidin. **(C)** Comparison of specific cytotoxicity induced by anti-HIV-1 sdAb CARs. **(D)** Comparison of specific cytotoxicity induced by MPER-targeting anti-HIV-1 CARs. Data represent the mean ± standard deviation from triplicate wells.

## Discussion

In recent years, CAR technology has been extensively expanded as an HIV-1 curative therapy [13, 15, 16, 26, 27, 47, 61-63]. Various anti-HIV-1 CARs targeting multiple different epitopes have been designed and evaluated their effector activities in in vitro and in vivo systems (See review[47]). However the different assay systems easily generate conflicting conclusions even using the same source of antibody for CAR design [20, 27, 47, 64, 65]. To obtain highly reproducible result, we first selected single cell clones which can stably express HIV-1 envelope proteins over 7 days upon DOX removal from the culture medium, named HXBC2 #39. We eliminated HLA surface expression and minimized levels of CD4 surface expression, resulting in low levels of CAR-independent cytotoxicity mediated by mismatched HLA or CD4-dependent syncytium formation. As a result, this cell enabled assessment of HIV-1 specific effector activity even at low E:T ratios (**Fig 3E**). The combination with accurately countable multicolor flow cytometry allowed us to provide quantitative and reproducible results with wide-dynamic range and low background. Importantly, any residual background mediated by CAR-independent cytotoxicities can easily be compensated by using an appropriate range of E:T ratios. With this assay, we successfully ranked 9 newly designed anti-HIV-1 CARs targeting various regions of HIV-1 envelope proteins. Three CARs targeting the MPER of gp41 showed approximately 2-fold higher activity than CARs targeting other regions of HIV-1 envelope proteins (**Figs 5 and 6**). Importantly, these results were obtained from two different CAR backbones constructed by scFv (10E8) or sdAb (2E7 & 2H10), suggesting that the MPER can be a good target for designing anti-HIV-1 CAR or other anti-HIV-1 biologics such as HIV-1 specific antibodies with effector activities.

In general, bNAbs targeting MPER are known to be more effective against broader range of HIV-1 strains but less effective and also display autoreactivity or polyreactivity [66, 67]. A similar observation was confirmed when the scFv form of 10E8 was tested for staining of HIV-1_HXBC2_ envelope proteins expressed on HCBX2#39 cells (**Fig 4B**). Whereas we did not observe such negative features with our three MPER CARs; they showed more potent anti-HIV-1 effector activity than other CARs. Such difference may be caused by the different mechanisms of action between neutralization and effector activity mediated by CAR format.

Our assay is suitable to evaluate anti-HIV-1 effector activity against HIV-1_HXBC2_ infection, and further validation is required against other virus strains. The bNAbs used here for designing anti-HIV-1 CARs are known to be effective against broad spectrum of HIV-1, and we expect that these CARs selected via our assay would be effective against other strains. Whereas one type of CAR would not be able to cover all HIV-1 species, the combination of two or three different anti-HIV-1 CAR molecules may be necessary to cover HIV-1 quasispecies within patients as used with the treatment by bNAbs. Since hundreds of anti-HIV-1 bNAbs are now been available for designing new CARs [68], our novel assay should be very powerful for selection of functional CAR molecules which cannot be identified by binding feature to epitope or neutralization activity. For further efficient CAR screening, it would be important to design additional target cells expressing more variety of HIV-1 envelope proteins, for example a CCR5 tropic HIV-1 strain. By marking other target cells with different fluorescent protein, we can assess multiple anti-HIV-1 CARs at the same time in the same assay system. CRISPR/Cas9 technology would be useful to modify the envelope sequence in HXBC2 #39 cells. It is also possible to generate patient specific target cells with the same system. As such, our assay is able to proceed developing further effective anti-HIV-1 CARs even in a tailor made fashion.

## Materials and Methods

### Cell preparation and culture

Jurkat cells inducibly expressing envelope of HIV-1 _HXBC2_ (NIH AIDS Reagent: #3953) by the removal of doxycycline (DOX) from culture medium were used as a parental cell for an HIV-1 specific target cell (designated as HXBC2). Jurkat cells missing the envelope expression were used as a parental cell for non-target control (AIDS Reagent: #3954, designated as ΔKS). The HLA-Class I expression of these cells were first eliminated by gene targeting of human-β2 microglobulin using CRISPR/Cas9 technology [69]. These cells were then genetically marked by fluorescent proteins, mCherry and TagBFP, respectively, to allow counting of absolute cell numbers of each populations by flow cytometry. HXBC2 cells were gene marked by two red fluorescent proteins for superior detection on MACSQuant analyzer 10 or MACSQuant VYB (MiltenyiBiotec). The cells were stained by anti-HLA-A,B,C antibody (W6/32, Biolegend) and anti-human CD4 antibody (RPA-T4, Biolegend), and populations with the double negative for HLA-A,B,C and CD4, but with the positive for mCherry or TagBFP were sorted out by the BD FACSAria II (BD Biosciences). Bulk sorted populations were further separated into single clone populations by a limiting dilution. Each clone was tested in an assay for induction of HIV-1_HXBC2_ envelope expression as well as CD4ζ CAR-induced cytotoxicity to confirm specificity and sensitivity for CAR-T inducing cytotoxicity. Absolute counts of the target and non-target cell populations were taken by MACSQuant and used to calculate relative cytotoxicity.

HXBC2 and ΔKS cells were maintained in Iscove’s Modified Dulbecco’s Medium (IMDM) (Invitrogen) supplemented with 15% fetal bovine serum (FBS) (Omega Scientific), Antibiotic-Antimycotic (ThermoFisher Scientific), Glutamax (ThermoFisher Scientific), 100 µg/ml of Hygromycin B (ThermoFisher Scientific), 100 µg/ml of Geneticin® (ThermoFisher Scientific), and 1 µg/ml of DOX (D3072, Sigma-Aldrich). All cells were incubated at 37°C and 5% CO_2_.

A lentiviral vector encoding anti-HIV-1 CAR was transduced and expressed in the enriched total human T cells obtained from fresh peripheral blood mononuclear cells by negative selection (EasySep Human T cell isolation kit, Stemcell Technologies). CAR-T cells were maintained in IMDM supplemented with 20% FBS, 30 IU/ml IL-2 (R&D Systems), Glutamax, and Antibiotic-Antimycotic.

### Plasmid construction

All vector plasmids were constructed by modifying the FG12 vector [55, 70, 71]. For cell labeling with a fluorescent protein, the cDNAs encoding mCherry [72] or TagBFP [73] were chemically synthesized and cloned into the FG12-based vector under EF-1 α promoter, respectively (GenScript). Fusion inhibitors, C46 [52, 53], V2o[58], and AP3[59] were also chemically synthesized based on the published sequences and cloned into the FG12 vector. A CRISPR/Cas9 lentiviral vector against human-β2 microglobulin was constructed by cloning an expression cassette for both Cas9 and guide RNA (gRNA) of PX458 (Addgene #48138) into the FG12 vector. The target sequence of gRNA was 5’-GAGTAGCGCGAGCACAGCTA-3’. ScFv CARs joined with a 41BB and CD3ζ chain at the C-terminus using human IgG4 Fc portion as a spacer, were inserted into the FG12 vector by swapping with the sequence for CD4ζ-P2A-EGFP [55, 74]. All CAR vectors contain two anti-HIV-1 genes, shRNA against HIV-1-LTR R region to protect CAR-T cells from HIV-1 infection [54, 55, 74] and C46 fusion inhibitor.

### Viruses

All lentivirus vectors were produced in 293T cells using calcium phosphate– mediated transient transfection with a packing plasmid (pMDGL), the pRSV-Rev, and the pCMV-VSV-G envelope protein plasmid as previously described [55, 70, 71]. The integration-defective CRISPR/Cas9 vectors were produced using D64E packing plasmid [75]. HIV-1_AD8_ and HIV-1_BaL_ stocks were prepared and infected to human primary CD4+ T cells as previously described [76].

### Flow-based cytotoxicity assay

Assay details were summarized in **Fig 2A**. Briefly, each HXBC2 and ΔKS cell was seeded in the same well of round bottom 96-well-plates (#3879, Corning Costar) at a density of 10,000 cells/well in 100 µl of IMDM containing 20% FBS. Effector-T cells modified by anti-HIV-1 CAR or a control vector with different Effector-to-Target ratios (E:T) in the same volume of medium were added to the well and incubated at 37°C (total 200 µl/well). Fifty µl of culture was taken following incubation and fixed in the same volume of 2% formaldehyde/PBS. Absolute cell numbers from each population were analyzed on a MACSQuant Analyzer 10 (Miltenyi Biotec Inc.) using FlowJo (AshLand). Relative cytotoxicity was calculated as a percentage defined by the equation: Relative cytotoxicity =100 x (1 – target cell number / non target cell number).

### SyAM production

All the SyAM constructs were chemically synthesized based upon the public database information available from Los Alamos National Laboratory (https://www.hiv.lanl.gov/components/sequence/HIV/neutralization/index.html) and cloned into the FG12 lentiviral vector. SyAM expressing lentiviral vector was transduced to 293T cells at MOI 5. Cells were then cultured in IMDM supplemented with 7% ultra-low IgG FBS (FB-06, Omega Scientific), Antibiotic-Antimycotic, and Glutamax for 4 days. Recombinant forms of SyAM in culture supernatant were isolated by protein A affinity resins (MabCapture, ThermoFisher Scientific) using Pierce gentle Ag/Ab elution buffer (ThermoFisher Scientific). After dialysis to PBS, SyAMs were concentrated approximately 1 mg/ml with Amicon Ultra Centrifugal Filters (MWCO 30kDa, ThermoFisher Scientific) and stored in -80 °C freezer. Integrity and purify of SyAMs were assessed by SDS-PAGE analysis using 8-16 % gradient gel (Lonza).

### Western blot

ΔKS and HXBC2 cells were cultured without DOX for 4 days to induce HIV-1 envelope protein expression. The same number of cells (2 x 10^6^ cells) were lysed in 100 µl of 1% CHAPS/PBS and analyzed on 4-20% SDS-PAGE gel (Lonza). The envelope proteins were detected by 2F5 (specific for gp41, NIH reagent #1475), 2G12 (specific for gp120, NIH reagent #1476), and horseradish peroxidase–conjugated antibody specific for human IgG (Santa Cruz Biotechnology, Dallas) as described previously [77].

### Immunofluorescent staining

Cell surface expression of HIV-1_HXBC2_ envelope proteins were detected as follows. One million HXBC2 #39 cells cultured in the absence of DOX for 10 days were incubated with recombinant sCD4 or SyAMs (1 µg each) on ice for 30 min, followed by Alexa488- or APC-conjugated anti-human Fc antibody (Jackson ImmunoResearch Laboratories Inc.). The expression of each anti-HIV-1 CAR was detected by protein L-biotin (GenScript) and Alexa488-conjugated anti-human Fc antibody or APC-conjugated anti-human Fc antibody as described elsewhere [78]. Antibodies for staining of intracellular cytokines, IFN-γ, TNF-α, and IL2, as well as human CD8 were purchased from Biolegend.

### HIV-1 specific T-cell proliferation assay

CAR-T cell proliferation assay was performed as follows. HXBC2 #39 or ΔKS #13 cells were cultured in the presence or absence of DOX for 10 days and co-cultured with CD4ζ CAR-T cells pre-labeled with cell proliferation dye eFluor 670 according to the manufacture’s instruction (ThermoFisher Scientific) in a 24-well plate over 6 days at E:T ratio = 1:10. Levels of T-cell proliferation in eFluor 670 positive population were monitored by a dye dilution assay using BD Fortessa and analyzed by FlowJo.

### Statistical analyses

Results are expressed as mean ± standard deviations (SDs). Errors depict SD. Statistical significance is presented with a *p*-value calculated via GraphPad Prism. The significance of survival-curve data was compared with a log-rank test. All other significance comparisons between groups were calculated by one-tailed unpaired *t*-test with Welch’s correction.

## Acknowledgments

We thank Jeffrey Brand for editing the manuscript.

## Supporting information

**S1 Fig.**
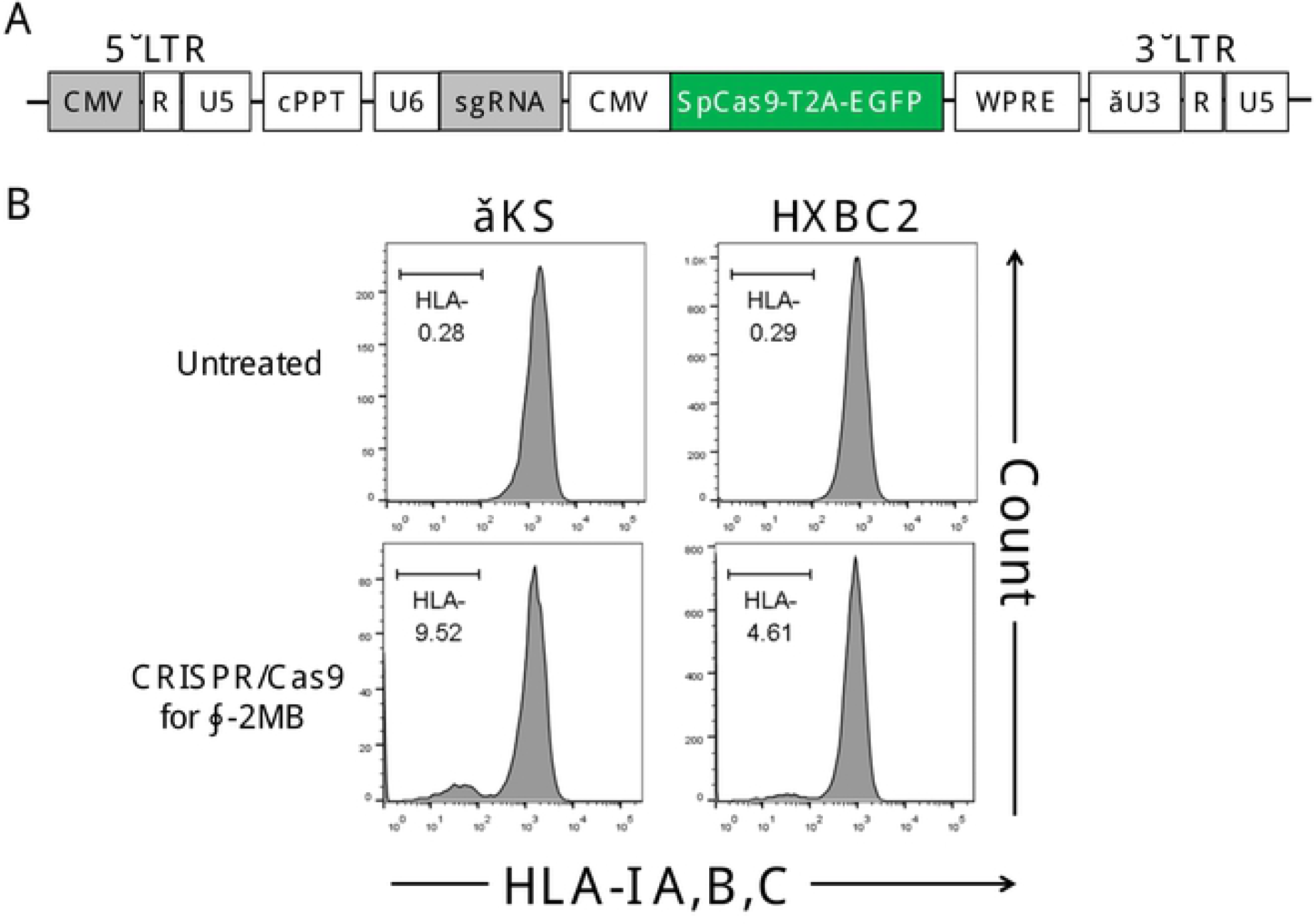
CRISPR-mediated knockout of HLA-I A,B,C surface expression via gene-editing of human-β2 microglobulin. **(A)** Schematic of a lentiviral vector encoding Cas9 together under CMV promoter (CMV) with single guide RNA (sgRNA). The vector has an FG12-derived backbone possessing a self-inactivating LTR, a central polypurine tract (cPPT), and a mutant Woodchuck Hepatitis Virus Posttranscriptional Regulatory Element (WPRE). T2A: self-cleaving 2A peptide of *Thosea asigna*. EGFP fused with Cas9 via T2A peptide (SpCas9-T2A-EGFP) serves as a transduction marker. sgRNA: specific for human-β2 microglobulin (β2MG) (5’-GAGTAGCGCGAGCACAGCTA-3’) under the human U6 RNA pol III promoter (U6). **(B)** Jurkat cells without (ΔKS) or with inducibly expressing HIV-1_HXBC2_ envelope proteins (HXBC2) were transduced with a non-integrating lentiviral vector encoding Cas9 together with β2MG sgRNA, which contain integrase with a D64E mutation. These cells were stained with PE-conjugated anti-human HLA-I A,B,C antibody.

**S2 Fig.**
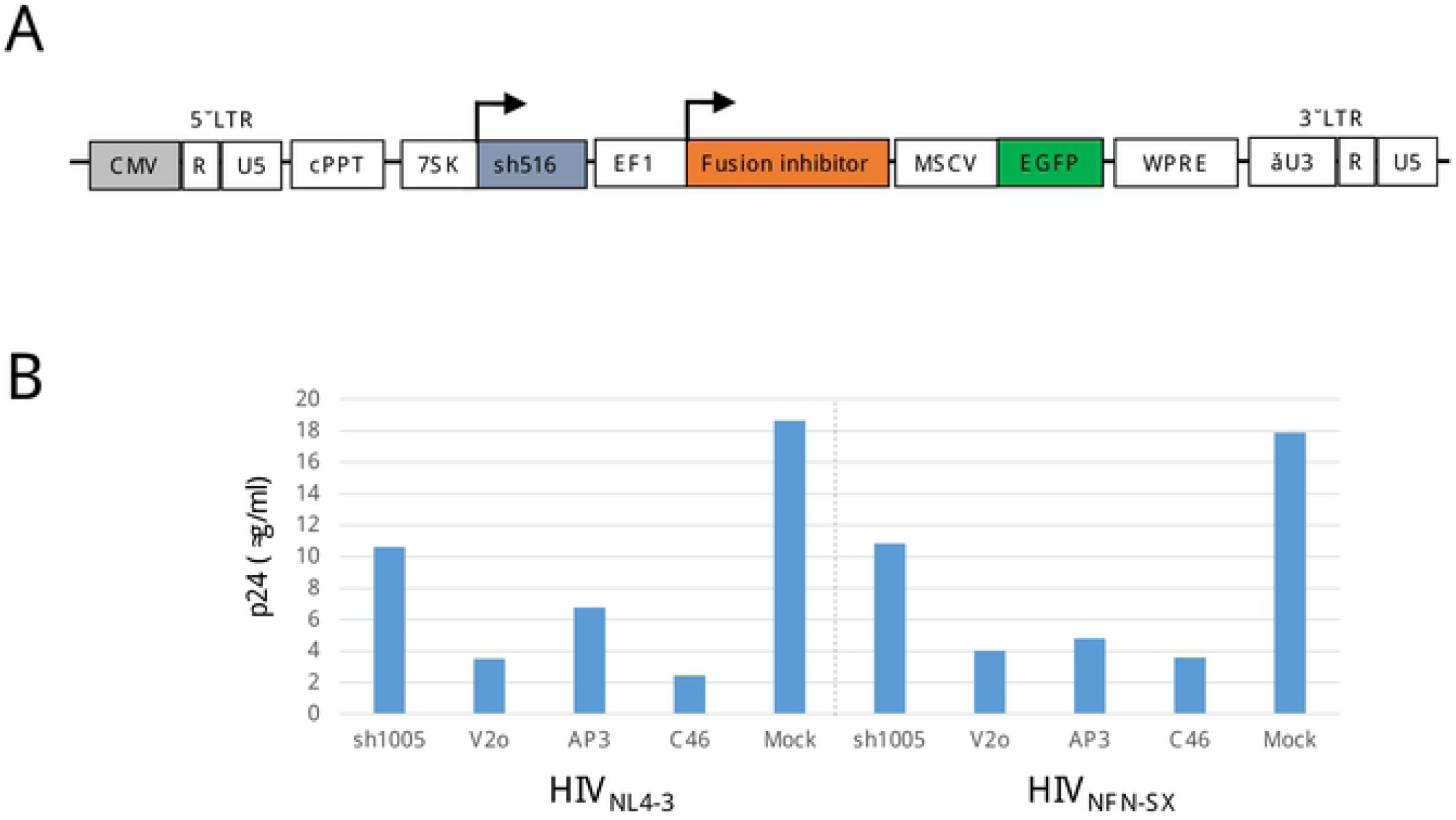
Comparison of anti-HIV-1 gene products against HIV-1 infections. **(A)** Schematic of a lentiviral vector encoding anti-HIV-1 chimeric antigen receptor (CAR) under the murine stem cell virus promoter (MSCV). The vector has an FG12-derived backbone possessing a self-inactivating LTR, a central polypurine tract (cPPT), and a mutant Woodchuck Hepatitis Virus Posttranscriptional Regulatory Element (WPRE). Dual: the vector encoding EGFP together with two anti-HIV-1 genes, sh516 targeting HIV-1 LTR R region under 7SK promoter (7SK) [76] and three different fusion inhibitors (V2o, AP3, or C46) under EF1-α promoter (EF1α) or sh1005 under H1 promoter[56]. **(B)** Human primary CD4+ T cells were positively selected by anti-human CD4 magnetic beads. Cells were stimulated by anti-CD3/CD28 antibodies (1 µg/ml each) for 24 hours in the presence of 30 IU/ml of human IL-2 and transduced with above vectors. EGFP-positive populations were enriched by FACS Aria II on day 3 post-vector transduction and infected by two different HIV-1 strains—HIV-1_NL4-3_ or HIV-1_NFN-SX_—at 100ng of HIV-1 p24 per one million cells. HIV-1 p24 amounts in culture supernatant were titrated 6 days post-HIV-1 challenge. Mock: non-lentiviral vector transduced cells.

